# Genome-Wide Search for Candidate Drivers of Adaptation Reveals Genes Enriched for Shifts in Purifying Selection (SPurS)

**DOI:** 10.1101/2020.01.11.902759

**Authors:** AB Popejoy, DE Domanska, JH Thomas

**Author notes:** Corresponding Author: AB Popejoy.

## Abstract

An open question in comparative evolutionary genomics is whether or not certain loci are the primary drivers of divergence between taxonomic lineages or species groups. Alternatively, genetic drivers of species divergence may be evenly distributed across the genome. The increasing availability of genome sequences from diverse taxa has enabled the development of novel methods to address this question. Genomes of many highly diverged species may now be compared in order to tease apart genetic differences that drive adaptive or functional divergence, and genetic differences that are observed by chance and are not causally linked to traits that differ between species or lineages. In order to test the hypothesis that a particular subset of loci or genes is responsible for driving adaptive changes between mammals and non-mammals, we developed a novel comparative approach to identify sites that are highly conserved within lineages or species groups and diverge between them. Loci with a high concentration of these sites may be called Shifts in Purifying Selection (SPurS) because a change has occurred between two groups of species at some point in the past, and the shift is conserved (via purifying selection) over a long period of time. Evaluating 7484 orthologous gene copies from 76 vertebrate species, we developed an empirical distribution of SPurS across the genome between Synapsida (placental and non-placental mammals) and Sauropsida (birds, crocodilians, squamates, and turtles), and compared this distribution to the expected null distribution of SPurS using matched simulated data. We then identified a subset of genes that is enriched for SPurS, relative to the full set of genes and to their matched simulated alignments. These SPurS-enriched genes are thus likely candidate drivers of functional divergence or adaptation between the mammalian and non-mammalian species groups in our analysis. Investigators seeking to identify genetic drivers of inter-species evolution may find this method useful, and we provide a web-based software interface to facilitate its use.

## Introduction

Identifying genetic variants that are responsible for observed phenotypic differences is a fundamental objective of modern genomics. In human genetics research, methods often compare individuals from different disease cohorts, ethnogeographic populations, and regions to identify candidate loci driving functional differences and population-level adaptive traits. Until recently, there has been less of a focus on inter-species comparisons due to the limited number of species with sequenced and published genomes. The construction of thousands of new genome assemblies from diverse species now provides an opportunity to develop inter-species approaches for identifying genes and variants that may be responsible for deeply rooted phenotypic differences, which are likely involved in driving functional adaptation between large groups of species sharing certain traits and/or taxonomic lineages.

A multitude of intra- and interspecific methods designed to detect signals of natural selection are well characterized in the literature (1). Purifying (or negative) selection eliminates mutations as they arise in a population, conserving the existing or ancestral state of a particular site in the genome. Methods that rely on purifying selection to elucidate potentially important genomic loci identify regions that are highly conserved in a comparative analysis. In inter-species analyses, these loci include ultra-conserved elements (UCEs) among distantly related species (2) or regions that are conserved within a taxonomic group such as mammals (3) or vertebrates (4).

In contrast to purifying selection, *positive* selection is the rise in frequency of new mutations that are evolutionarily advantageous in certain contexts, which often leads to many site-specific differences between species. Sites that have undergone positive selection in multiple species or lineages are considered drivers of adaptation, and common methods to detect these recurrent positive selection events involve the detection of multiple different alleles across species (5). Approaches that detect these sites, where recurrent positive selection events have occurred, focus on the identification of loci that are flexibly adaptive across different species, with different amino acids providing various selective advantages for individual or closely related taxa.

One limitation of focusing on genetic loci that are either highly conserved (via purifying selection) or highly divergent (via positive selection) is that these approaches do not account for loci that are divergent between specific groups of taxa, and that are also highly conserved over long periods of time. For example, consider a single amino acid substitution that occurred in the most recent common ancestor (MRCA) of species in two very large and ancient taxonomic lineages, such as mammals and non-mammals. Perhaps this substitution was advantageous for the mammalian descendants of the ancestral population, while the original or ancestral residue remained beneficial for its non-mammalian descendants. In cases where such a change (or multiple changes) at a particular locus leads to functional differences between mammals and non-mammals, these cases would be considered genetic drivers of adaptation between the two species groups.

At these adaptive loci, a multiple alignment of nucleotide or amino acid sequences from various mammals and non-mammals would show all (or most) mammals having one residue, while all non-mammals would have another. Neither positive nor purifying selection-based methods would detect such a signal. Because a single residue is not highly conserved across most taxa in the alignment, it would not be considered under strong purifying selection, and because there is only a single change across all taxa, neither would it be considered under positive selection. In this paper, we describe a novel, sequence-based approach to detecting loci with this pattern of divergence and conservation, such that all species in one group favor one amino acid residue, while species in another group favor a different one.

## New Approach: Shifts in Purifying Selection (SPurS)

The method described in this paper is designed to evaluate sites in a multiple-species alignment and identify loci with many sites that exhibit only a single substitution event (or very few) across a large number of orthologous sequences from distantly related species. Conceptually, genomic loci may be considered candidate drivers of adaptation between two groups of species when they exhibit a high concentration of sites with only two conserved amino acid residues, each of which is present in all species of one group and none in the other. Sites in multiple alignments that show this pattern of amino acid segregation between two species groups (e.g. mammals and non-mammals) are called *Shifts in Purifying Selection* (SPurS) because the pattern is a result of a *shift* or change in the amino acid residue that is then conserved under purifying selection between two species groups over long periods of time.

This approach has several advantages over other methods for detecting candidate gene drivers of adaptation. Common inter-species methods to scan the genome for sites of positive selection lack power to detect a single impactful shift between lineages or species because they often require recurrent substitution events at the same site (6,7). These approaches are also limited to comparing species that are not too distantly related, such that the number of synonymous changes between them is not saturated (8). Positive selection scans may assume uniformity across loci and lineages (9), which presents a substantial limitation to identifying regions of the genome that differ between specific groups of species, regardless of their taxonomic relationship. Some rate- and model-based approaches to identifying regions of accelerated evolution have suggested the ability to elucidate parallel patterns of selection in different genes and lineages, and some methods directly account for heterogeneous evolutionary rates across sites and branches (10,11). However, these methods often operate on *a priori* assumptions or constraints about the phylogenetic relationships between species in the analysis and other parameters, which complicate the interpretation of findings.

SPurS can be calculated on any multiple-species alignment without setting any priors other than defining to which group each taxon belongs, and a signal may be found when comparing any two groups of species, regardless of their taxonomic or phylogenetic relationships. This method makes no assumption of rate uniformity across loci, lineages, or taxa. It is based on a heuristic statistical measurement at each site, so there are no model assumptions to complicate the interpretation. Loci identified as having shifts in purifying selection are simply protein alignments with a large relative proportion of sites where amino acid residues segregate in two different lineages or trait-defined groups of species.

Figure 1 illustrates the conceptual framework for SPurS, showing a single residue conserved in each group of species (Fig.1A) or taxonomic clade (Fig.1B). A “complete SPurS site” (Fig.1C) occurs when exactly two amino acid residues are found at a site in the alignment, and all members within a particular group or clade of species shares the same residue. Species groups can either be defined on the basis of their taxonomic relationship (i.e., clades in a phylogenetic tree), or on the basis of any other condition, such as a shared trait or phenotype (e.g., echolocation or the presence of wings). In the latter case, SPurS-enriched genes could be considered candidates for convergent evolution, since the species groups are defined on the basis of a common trait, independent of their evolutionary relationships. For the purpose of describing this method, which may be applied to groups of species defined on the basis of phenotypes or on phylogenetics, we refer to any SPurS comparison groups as “species groups” moving forward.

**Figure 1.**
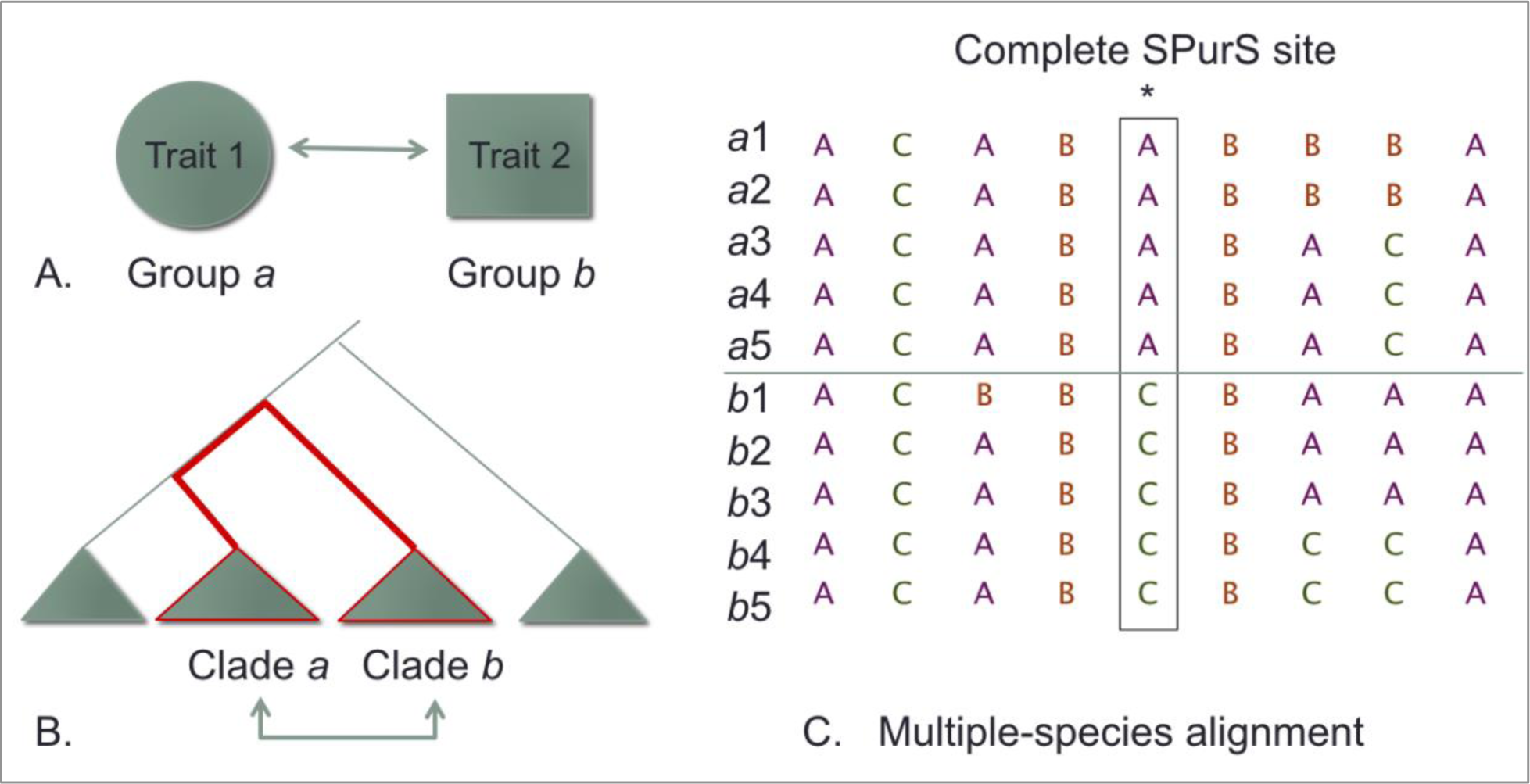
Conceptual Illustration of Shifts in Purifying Selection (SPurS). Species to be compared are classified into two categories, which may be pre-identified on the basis of trait differences (A), phylogenetic clade membership (B), or any other variable of interest. Protein sequences from all species from both groups are aligned (C) and each site in the alignment is evaluated individually.

### I. SPurS Algorithm

The concept of detecting within- and between-group differences in population genetics can be traced back to Sewall Wright’s F-statistics in the 1920s (FST) (12). The basic idea is to measure pairwise differences (heterozygosity) between individuals sampled from different populations to detect underlying structure. In this formulation of the SPurS statistic, we re-work the idea of pairwise heterozygosity between populations to measure the ratio of between- and within-group divergence across large lineages in a phylogeny or other categories of species. Our FST-like statistic designed to quantify shifts in purifying selection using Ψ (Psi), conveniently an acronym for <u>p</u>urifying <u>s</u>election shifts, and measures divergence between species from the same group, compared to divergence between species from different groups. Thus, Ψ is calculated as a difference of *between*-group divergence and *within*-group divergence:

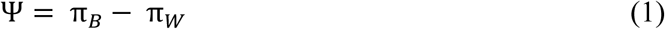

where π_*B*_ is the ratio of the total number of pairwise differences in residues *between* species in different groups G1 and G2 to the total number of comparisons made between groups,

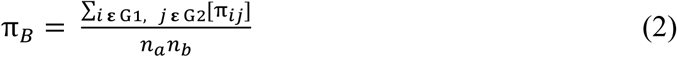

and π_*W*_ is the ratio of the total number of pairwise differences among species *within* the same group, compared to the total number of comparisons made within groups,

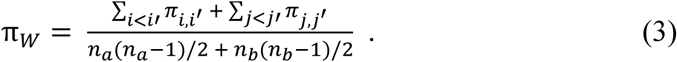

In both (2) and (3), *n*_*a*_ and *n*_*b*_ are the numbers of species analyzed from groups 1 and 2 (G1 and G2), respectively. These are not always equal to the number of species from each group that are present in the multiple alignment; if a residue for a particular species is not present at the site being analyzed (a gap in the alignment) then this taxon is not counted toward *n* in the relevant group.

In equation (2), π_*ij*_ represents a comparison of residues at site *S* between species *i* and *j*, one from each of two different species groups. This is done for all possible pairings of species in different groups. When the residues are the same, (*Si* = *Sj*), π_*ij*_ = 0 and and when they are different (*Si* ≠ *Sj*), π_*ij*_ = 1. Thus, when ∑π_*ij*_ = 0, there is no divergence between species from different groups (π_*B*_ = 0). When the number of divergent sites equals the total number of comparisons made, ∑π_*ij*_ = n_*a*_ ∗ *n*_*b*_, then π_*B*_ = 1, indicating complete divergence between groups at this site, since all residues compared between groups are different from one another.

In equation (3), *π*_*i,i*_′ represents a pairwise comparison of residues at site *S* from two species *i* and *i’* both sampled from the same species group (e.g. G1), and *π*_*j,j*_′ represents a pairwise comparison of residues at site *S* from two species *j* and *j’* sampled from G2. Again, this is done for all possible pairs of species in the same group. As above, when the residues compared between two species are the same (*Si* = *Si’* or *Sj* = *Sj’*), there is no divergence between them; thus π_*i,i*_′ = 0 or π_*j,j*_′ = 0. If the residues between species are different (*Si* ≠ *Si’* or *Sj* ≠ *Sj’*), then *π*_*i,i*_′ = 1 or *π*_*j,j*_′ = 1. If the number of divergent residues within groups 1 and 2 at site *S* is equal to the total number of pairwise comparisons among species within groups, then

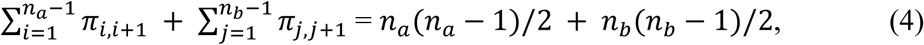

and π_*W*_ = 1, indicating that all residues from species within each group are divergent from one another. The case of greatest interest, indicating a “complete” shift in purifying selection, is one in which all species diverge between groups, and none diverge within groups. That is, when:

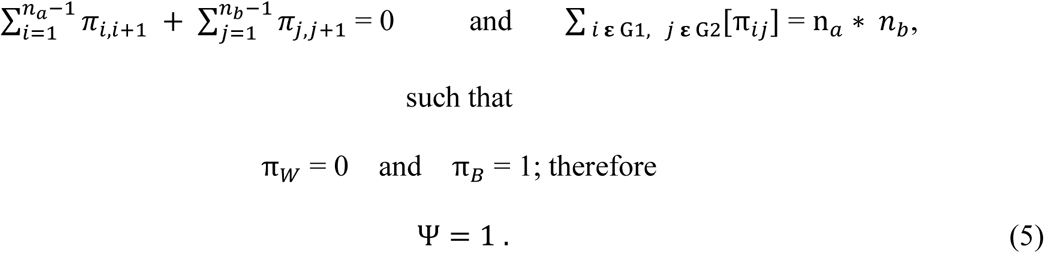

### II. Interpreting Ψ (Psi)

Cases (1) through (4) in Fig. 2 demonstrate how to interpret Ψ values. When Ψ < 0, amino acid residues differ more frequently between species in the same group than species in different groups. This denotes an excess of within-group divergence. Case (2) describes sites at which there is equal within- and between-group divergence (Ψ = 0). This situation arises when there is any amount of divergence or conservation in the alignment, so long as it is equivalent between species groups. In some of these cases, there is complete conservation at a site, indicating some mixture of strong purifying selection or insufficient divergence time for a change to occur. Case (3) describes sites at which Ψ > 0, indicating more divergence between groups than within them. Case (4) refers to “complete SPurS sites” in which Ψ = 1. In this case, the site is characterized by a single change or substitution, followed by conservation of the two group-specific states over a long period of time.

**Figure 2.**
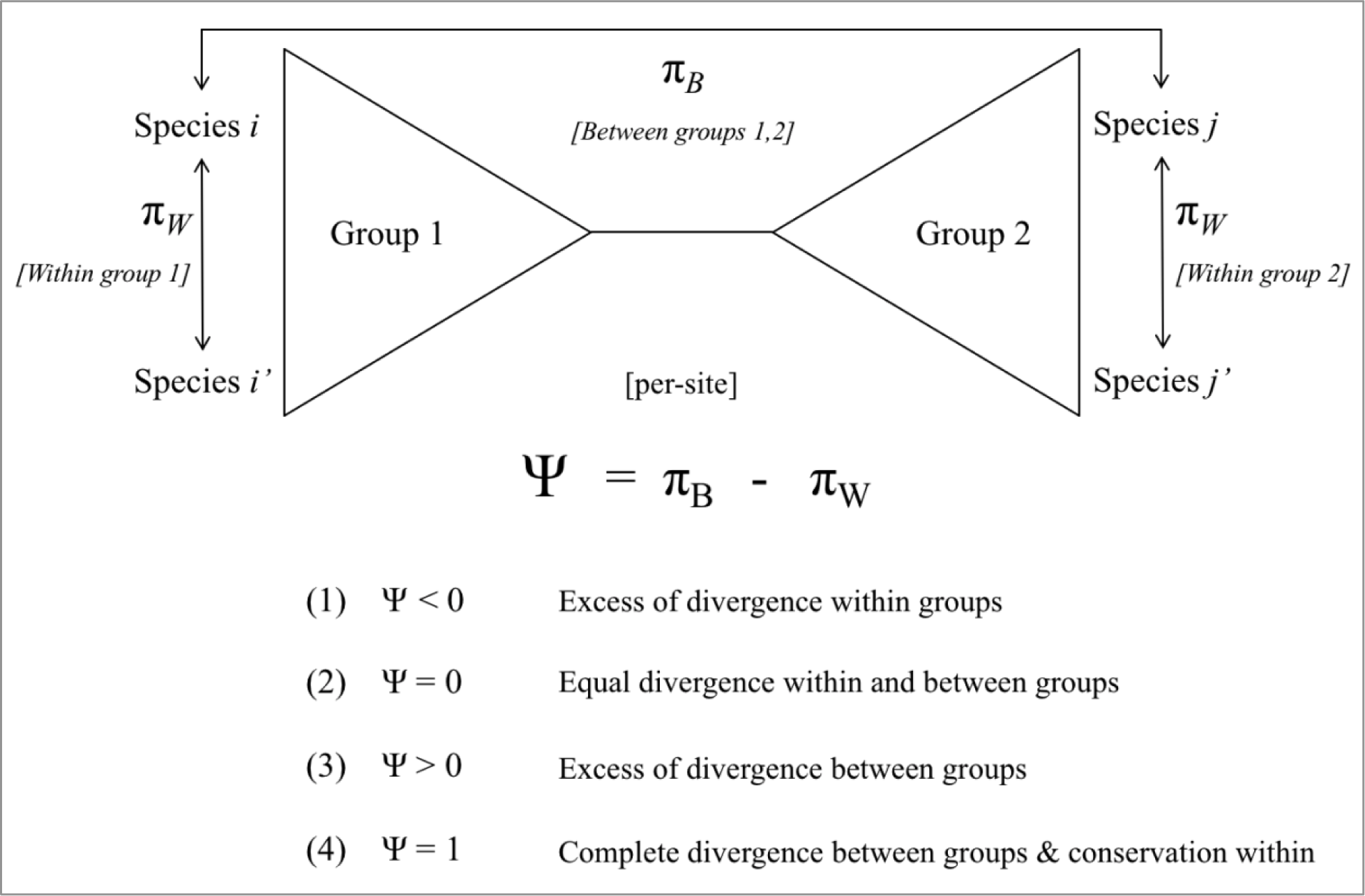
Diagram and Statistical Properties of Ψ. In this simplified illustration of Ψ, Groups 1 and 2 represent species to be compared; πB represents the number of differences in residues at a single site among all pairwise comparisons of species in different groups (normalized by the total number of comparisons), and πW is the total number of differences at the same site among species within each group, normalized by the number of pairwise within-group comparisons. Ψ value interpretations are shown in cases (1) through (4), where each case represents a combination of divergence (π) or conservation among species both within and between species groups.

### III. Genome-Wide Test for SPurS

The purpose of this study is to determine whether patterns of genetic difference between species groups and conservation within groups is arbitrary, or whether these patterns are indicative of an underlying evolutionary mechanism for divergence between species groups. Our null hypothesis is that SPurS are 1) observed by chance, and 2) randomly distributed across the genome. If the null hypothesis holds, then the pattern we have described occurs by chance. Our alternative hypothesis is that 1) SPurS are not random, but rather occur more or less often than would be expected, and 2) a subset of genes is responsible for species group divergence and thus should exhibit a higher concentration of SPurS relative to other genes.

In order to test this hypothesis, we designed a validation study to identify SPurS-like patterns across the genome using the algorithm previously described. This ‘genome-wide’ test for SPurS includes calculating Ψ across all sites and comparing those values to matched simulated data. It also includes calculating average Ψ-values within genes and comparing the concentration of SPurS patterns in real and matched simulated alignments. The goal of this approach is to construct a conservative simulated “null” dataset to identify real genes enriched for SPurS sites, relative to simulated data. Since the simulated data generator implements uniform purifying selection across the whole tree (reflected in the amino acid change rate matrix), comparing the real protein alignments to simulated data provides a direct test of both the frequency and concentration of observed SPurS across loci.

## Materials and Methods

### I. Selection of Species Groups and Individual Taxa

To conduct this genome-wide test of the SPurS algorithm, we chose to define comparison groups of species corresponding to two major taxonomic clades that are roughly equivalent to mammals and non-mammals. By sampling the evolutionary tree of vertebrates with fully sequenced genomes, we selected 76 species representing the two large phylogenetic clades with long internal branch lengths: 44 from Synapsida (monotremes, marsupials, and placental mammals) and 32 from Sauropsida (birds, crocodilians, squamates, and turtles). The inclusion of species that are distantly related *within* groups is critical, in order to minimize the random occurrence of SPurS signals. This is because species that are closely related share the vast majority of identical sites in the genome. Figure S-1 and Table S-1 (Supplementary Materials) illustrate the phylogenetic species tree and list of scientific names, respectively, for all taxa included in the analysis.

### II. Identification and Alignment of Orthologous Genes

All sequence and species data were obtained from Ensembl (13), BGI (14), or NCBI RefSeq (15). We used genes identified as orthologues in mammals by OrthoMam (16) and identified as 1:1 orthologues between human and coelacanth (17) to identify probable 1:1 orthologues throughout Amniotes. In 22 cases, a probable paralogue (duplicated and diverged version of the ancestral orthologue) was present in Sauropsida and these genes were removed from analysis. Future studies should pay close attention to paralogues, since two-copy paralogues that are reciprocally lost in the clades under study often produce a strong SPurS signal. 7484 orthologous genes were identified as being shared among a majority of the species (>55) included in the analysis. Alignments were generated using orthologous protein sequences obtained through BLAST search capture (18) and the MUSCLE protein alignment tool (19). No manual adjustments were made to the alignments.

### III. Empirical Distribution of Ψ Across Sites and Within Genes

The SPurS heuristic **Ψ** was calculated independently for each column of 4.7 million aligned amino acid residues (“sites”) from all taxa sharing orthologous gene copies. Figure S-2 (Supplementary Materials) shows the number of taxa that have an orthologous sequence for each gene. Only sites where a sufficient number of species (>50%) have a conserved residue were evaluated.

In addition to calculating the overall distribution of **Ψ** across all sites, we calculated both the frequency of ‘complete SPurS sites’ (where Ψ = 1) in each orthologous protein alignment and the average Ψ value across each alignment, in order to determine whether there are trends in Ψ values observed in some subset of genes, relative to other genes and compared with simulated data.

### IV. Matched Simulated Data with Evolutionary Rate Multiplier

Simulated data were generated assuming a model of uniform purifying selection and conditioned on the evolutionary relationships among the species included in the analysis, then tailored to match each unique protein on sequence length as well as the number and taxa present in the alignment. This dataset represents what we would expect to observe if the evolutionary process driving patterns of similarities and differences were only dependent on a general model of amino acid substitution and phylogenetic relationships among species, assuming rate uniformity across branches and amino acid sites. The process involved an initial simulation of 3 million sites of a multiple-species alignment under the JTT model of amino acid substitution, which sets the relative rates of amino acid substitutions (20). The simulation was conducted in Seq-Gen (21) and conditioned on the species phylogeny of taxa included in the real data (Fig. S-1).

In order to be conservative in our estimate of the null distribution, we also determined the evolutionary rate multiplier that would maximize the proportion of complete SPurS sites and used this multiplier to conduct the simulation. Figure S-3 (Supplementary Materials) shows the proportion of complete SPurS sites for simulations (N=38) that were conducted using a range of multipliers [0.01-2.0] to identify the appropriate rate for constructing a conservative null dataset.

The initial alignment of 3M sites was broken up into individual alignments, each matching the length of one real protein, and trimmed to include sequences for only those species with an orthologous sequence for that protein. The process of matching each simulated alignment on length and exact species represented to an alignment in the real dataset allows us to simulate the number of SPurS that would be observed by chance, given the evolutionary relationships among species. This enables us to directly compare each real protein alignment to a matched simulated alignment, while also providing an analogous comparison between the real and simulated datasets as a whole.

### V. Comparison of SPurS in Real Alignments to Simulated Dataset

In order to test whether a SPurS-like pattern is observed in multiple alignments more often than would be expected by chance, the distribution of Ψ across all sites among simulated data was compared to the Ψ distribution across sites in real proteins. In the absence of selective pressures that have preserved these sites over long periods of time, the distribution of Ψ (indicating the frequency of observed SPurS patterns) should not differ between the real and simulated alignments. Therefore, comparing the distribution of Ψ between these two datasets provides a natural test of the null hypothesis that SPurS occur as often as expected by chance. Because simulated data were conditioned on the species phylogeny of taxa included in our analysis (Fig. S-1) and matched to the length and number of taxa included in each real alignment of orthologous genes (Fig. S-2), the comparison of real and simulated data constitutes a test of whether SPurS occur more or less often than would be expected under the null hypothesis. This is also a conservative comparison, since the simulated data were generated with a rate parameter designed to maximize the random occurrence of complete SPurS sites.

In addition to comparing the overall distribution of Ψ across all sites in real and simulated alignments, we calculated the frequency of complete SPurS sites in each alignment from real sequence data and compared this to the frequency of complete SPurS sites in their matched simulated alignments. This comparison enables us to test whether certain genes are more likely to exhibit SPurS-like patterns by chance, or whether we would expect these patterns to be evenly distributed across genes under the null. If a subset of real alignments exhibits elevated SPurS signals relative to others, and no there is no such difference in the frequency of SPurS among equivalent simulated alignments, this would suggest that a true evolutionary mechanism is at play.

### VI. SPurS Analysis Toolkit on the Genomic Hyperbrowser

Considering the potential for applications of SPurS to a diverse array of biological questions, there may be researchers with minimal bioinformatics skills who would benefit from its use. As such, we have designed an online toolkit for researchers seeking to identify genomic loci that are likely involved in adaptive trait differences between large groups of species and have made it accessible to a wide range of users. The user-interface is available on the Genomic Hyperbrowser^1^ platform for genomic data analysis, hosted at the Institute for Bioinformatics at the University of Oslo. The Genomic Hyperbrowser is built and hosted on *Galaxy*, an open data-sharing platform designed for “accessible, reproducible, and transparent computational biomedical research” that allows users to freely access datasets and conduct analyses on their own datasets via an online user-interface (22).

The SPurS analysis toolkit features data visualization in the form of circular ‘SPurS plots’ along with histograms and table results, which allow for immediate visual comparisons of the distribution of SPurS across a protein with its simulated counterpart. All results, including ordered vectors of Ψ for both real and simulated data, are available for download on the results page. Users may select one gene at a time, and choose whether to compare mammals to birds, or mammals to Sauropsids (which includes birds). There is also an option to upload new alignments, as well as revised control files, so that users may conduct their own analyses on independent datasets or compare different subsets of species from our existing alignments. The SPurS program is also available in Python on A.B. Popejoy’s GitHub account, which details instructions on how to use the command-line functions of the program. Upon publication, all of the raw alignments from real and simulated data will be integrated into the Hyperbrowser for users to access and analyze.

## Results

### I. Distribution of **Ψ** in Real and Simulated Data

The ‘genome-wide’ distribution of Ψ is illustrated in Fig.3, with real data represented in purple and simulated data in gold. The purple and gold bars indicate the proportion of all sites in the analysis (y-axis) and their Ψ values, which correspond to discrete bins representing a range of Ψ values (x-axis). From looking at this figure, it is apparent that sites are not evenly distributed across the spectrum of Ψ values. Approximately 65% of all sites have Ψ < 0.1, and approximately 20% of sites have Ψ > 0.6. The remaining 15% fall between 0.1 and 0.6 and are evenly distributed.

**Figure 3.**
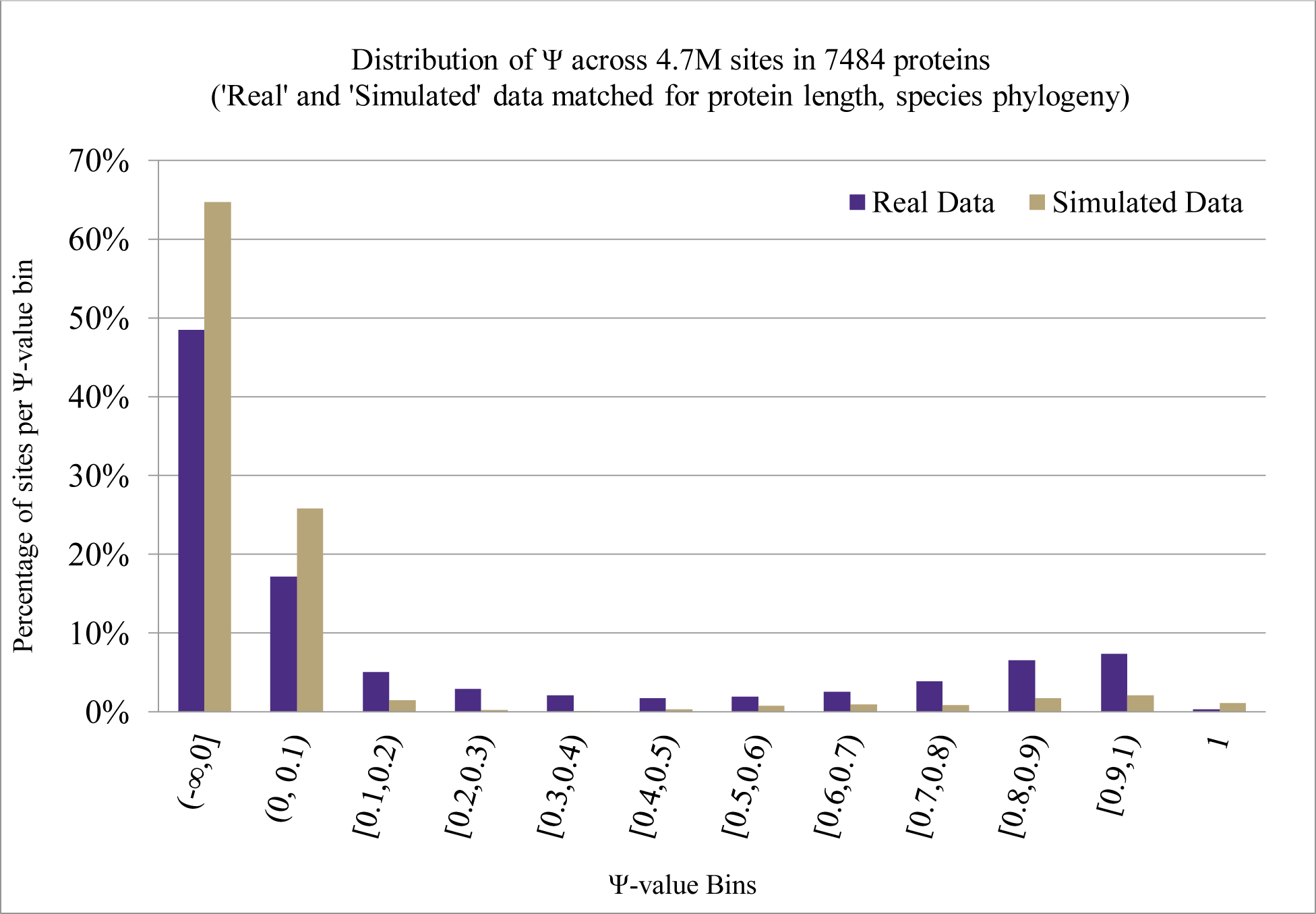
Genome-wide distribution of Ψ. Illustrates proportion of sites (y-axis) in each ordered categorical bin of Ψ value ranges (x-axis). Each simulated alignment was generated to empirically maximize the proportion of sites with Ψ=1 (rate multiplier = 0.2; See Fig. S-3 in Supplementary Materials) and matched to a real alignment on protein length and combination of taxa.

Despite the fact that the simulated data were explicitly conditioned to maximize the proportion of sites with Ψ = 1, neither real nor simulated datasets show a high proportion of complete SPurS sites. On the contrary, the majority of sites in both real and simulated data are in the lower range. Of these sites in the lower range, most have Ψ = 0. These sites are either highly conserved, and thus there are no within- or between-group differences observed; or the frequencies of within- and between-group differences are equivalent. Sites with Ψ < 0 exhibit greater divergence *within* species groups than between them, and the large proportion of these sites relative to those with higher Ψ values indicates long internal branch lengths of species groups on the phylogenetic tree.

Our results demonstrate a difference in the distribution of Ψ between real and simulated data; 90.5% of simulated sites and only 65.5% of real sites have Ψ < 0.1. Also, the distribution of real data is skewed right toward higher Ψ values, relative to simulated data, with the exception of the most extreme category (Ψ = 1). In this case, simulated data have a significantly higher frequency of complete SPurS sites (1.1%) compared with real data (0.32%) (*X*2 = 20250.08, p-value < 1.3e-3112, df = 1). What this shows is that our simulations were successful in maximizing the frequency of these sites, and they are exceedingly rare in both real and simulated data.

### II. SPurS are Concentrated in a Subset of Real Genes

Relative to matched simulated data and the full set of real alignments analyzed, a small subset of proteins had much greater concentrations of SPurS than expected. In some real protein alignments, 6.5% of sites exhibit a complete SPurS, whereas the vast majority of real data sites had fewer than 0.1% complete SPurS and no more than 1% of sites with this pattern was observed in any simulated alignment. The maximum percentage of complete SPurS sites for any simulated dataset was 0.84%, with a rate multiplier of 0.2 applied to the total branch length of the phylogenetic tree on which the simulations were conditioned.

Only 22 genes (out of 7484) have an average Ψ across sites > 0.6. Among these genes with high average Ψ values, several of them have a high frequency of sites with Ψ = 1. Since the maximum percentage of complete SPurS sites observed across simulated data was 1.1%, we suggest that any gene with a higher proportion of complete SPurS is worth following up for functional importance. The top 10 genes we sampled with the highest percentage of sites with Ψ=1 are: TMIE (6.5%), FAM81A (6.5%), ARVCF (6.5%), WNT2B (5.7%), GMPPA (5.5%), TMEM121 (5.1%), RHOV (5.1%), PACSIN3 (4.5%), CHAC1 (4.4%), OLFML3 (4.3%). A Gene Ontology analysis comparing these genes to all others in the dataset yielded no significant results.

### III. 3SPurS Plots

This ‘SPurS plot’ is a circularization of Ψ values at each site across the linear (sequential) protein alignment, and the first site in the sequence is always at the [12 o’clock] apex of the plot. Each line in the plot (referred to as ‘spokes’) represents the magnitude of Ψ, such that apparent gaps in spokes indicate low Ψ values (approximately 0) and long spokes in the plot indicate higher Ψ values (approximately 1). Uniformity of the spokes along the SPurS plot implies an even distribution of Ψ values across the protein, and long spokes clumped together in a plot indicate regions of SPurS signals with heterogeneity across sites in the protein.

Comparison of the SPurS plot for a particular protein to its matched simulated dataset is helpful in visually determining whether SPurS signals observed in any given gene are in fact suggestive of a shift in purifying selection, or whether this observation is due to an artifact of the particular species phylogeny of taxa included in the analysis.

Figure 4 illustrates the results of such a comparison, using SLC1A3 and FAM81A as examples. SLC1A3 shows no significant difference between the real and simulated datasets (*X*_2_ = 0.4, p-value > 0.8), despite several segments of long spokes (indicating high Ψ values), while FAM81A is significantly enriched for SPurS relative to its matched simulated alignment (*X*2 = 14.32, p-value < 0.0001). This comparison clearly demonstrates that some genes have a large concentration of sites with high Ψ values, including complete SPurS (Ψ = 1) while others have a roughly equivalent concentration of these sites compared to matched simulated data.

**Figure 4.**
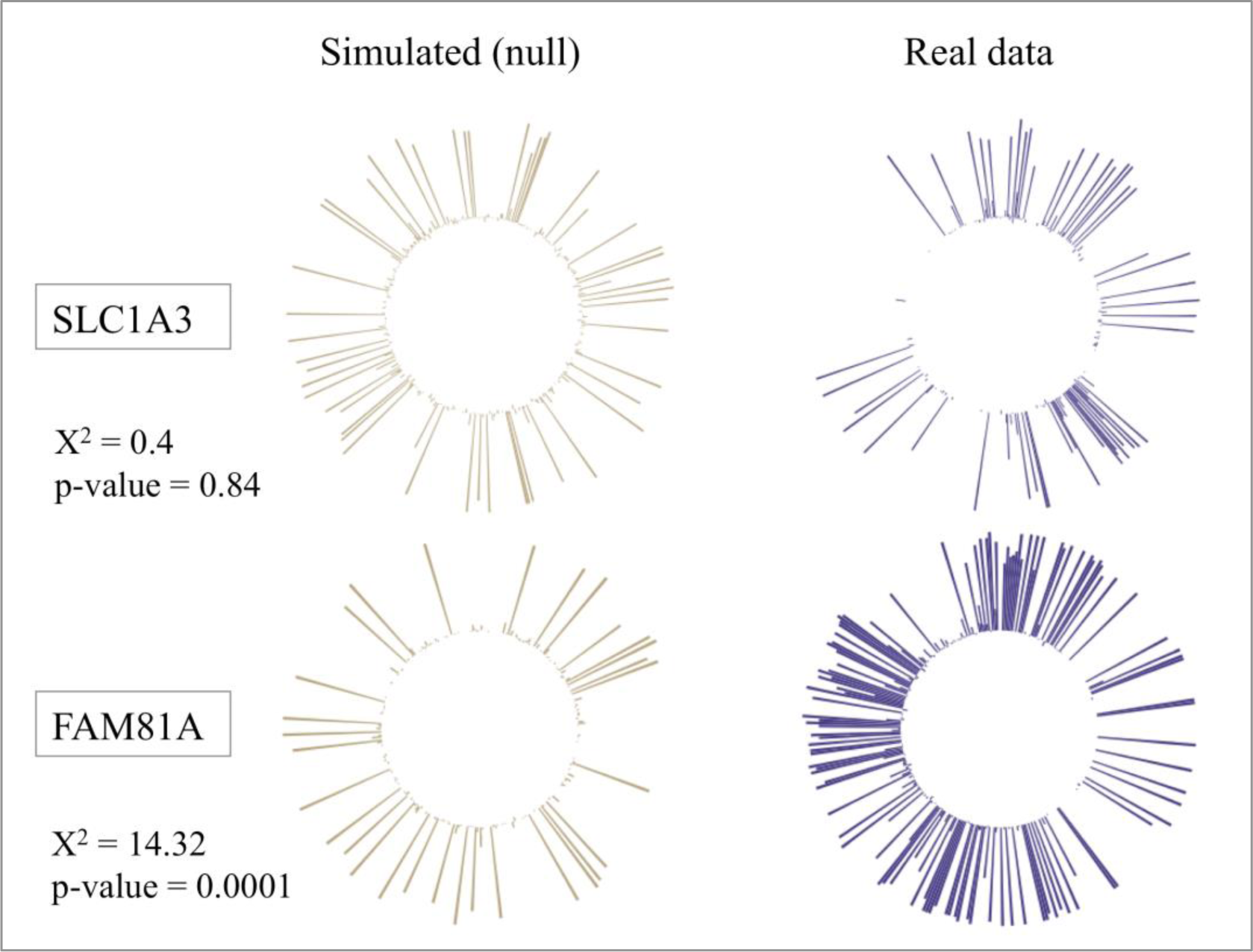
SPurS plots and statistical analysis results comparing mammals and birds (from the SPurS Toolkit on the Genomic Hyperbrowser). SPurS plots illustrate per-site Ψ along the physical sequences of two real proteins (right, purple) and their matched simulated datasets (left, gold).

## Discussion

SPurS-enriched genes can be considered candidate drivers of adaptation, likely contributing to phenotypic differences between pre-specified species groups or phylogenetic clades. In this case, our groups were specified according to taxonomic lineages, so the genes we have identified with a high frequency of complete SPurS sites are candidates for underlying phenotypic differences between Sauropsida and Synapsida. If the groups were specified according to a particular trait, as opposed to phylogenetic lineages, the SPurS methodology could be used to identify genes that are responsible for driving differences in that trait between species groups.

Our results demonstrate that although rare, SPurS-like patterns exist more often than expected under the null, and that sites with high Ψ values are concentrated in a subset of genes, rather than randomly distributed across the genome. Based on these results, we reject the null hypothesis that SPurS are observed by chance alone. We also reject the hypothesis that SPurS signals are evenly distributed across the genome, in favor of our alternative hypotheses that these patterns exist as a potential mechanism of evolutionary divergence between species groups, and that a subset of genes is primarily responsible for driving that mechanism. As this paper presents a proof-of-concept for a novel methodology, we consider this research to be hypothesis-generating. Thus, we believe that genes with a high concentration of SPurS sites are likely candidate drivers of divergence between species groups and suggest these findings be followed up with further inquiry to identify their functional mechanisms.

SPurS have likely been observed anecdotally by manual inspection of multiple-protein alignments from different groups of species; that is, amino acid sites with two different residues present between two large lineages or types of species such as mammals, birds, and fish. However, there has not (until now) been a general method to describe such sites and software provided open access to the research community in order to identify them systematically. We assume that the absence of a prior method to describe the SPurS phenomenon is most likely due to a dearth of available genomic sequences for species with an adequate amount of evolutionary distance between them. If species in one clade or group are too similar to one another (the total branch length among them is short), and the differences between clades or groups being compared is very distant (branch length connecting the two clades or between species groups is long), then one might expect to observe a large number of instances of SPurS by chance. For example, a comparison of several closely related primate species to several closely related fish might spuriously show a very high rate of SPurS sites, based on the evolutionary distance between the groups and evolutionary proximity within the groups. In our dataset, SPurS appear to be quite rare, indicating that branch lengths are sufficiently long within species groups, relative to the branch length between them.

We present a novel statistical method and software (both command-line and with an online user interface) to identify sites where SPurS occur. We provide access to all of our protein alignments, as well as the simulated datasets we used to compare to the real data, which enables others to replicate our findings and conduct original research with newly specified species groups. As such, we anticipate that the accessibility of our software and data, as well as the general nature of the SPurS detection methods, will open up a new area of research on genes that are candidates for drivers of adaptation between groups of species, either based on traits (detecting convergence) or on phylogenies (detecting genes responsible for species adaptation). In either case, we may be able to uncover phenotypes associated with SPurS-enriched genes for which there is no previously known function, by assessing the traits that differentiate the compared lineages or species groups.

The SPurS method can be applied broadly, equally relevant to other kinds of species beyond those in our analysis, including invertebrates and bacteria. Additionally, this method is general and adaptable to future improvements such as modifying it to identify shifts in purifying selection from nucleotide sequences in addition to protein sequences. This will enable an assessment of SPurS in non-coding regions of the genome, which may be an effective way to identify regulatory regions such as transcription factor binding-sites that differentiate between groups of species. There may also be further methodological developments to improve the method, including possible inclusion of maximum likelihood, and model-based, branch-site approaches. The full extent of future applications of this method have yet to be realized, and we predict that the increasing availability of data from diverse species will facilitate its use among computational and field biologists alike.

## Acknowledgements

Joseph Felsenstein served as the corresponding author’s PhD Supervisory Committee Chair during this project at the University of Washington. He provided thoughtful feedback on the mathematical framework and study design, specifically recommending comparison to simulated data constructed using an appropriate rate multiplier. Elizabeth Thompson was also a PhD Supervisory Committee Member and contributed comments to early conceptual framing of this study. Geir Kjetil at the University of Oslo approved the addition of the SPurS tool to the Genomic Hyperbrowser.

## Footnotes

1 https://hyperbrowser.uio.no/spurs/

2 All p-values for chi-square tests were calculated in Python, using various built-in functions. The initial function used was chi2.df from scipy.stats, which has an upper limit of 17 significant digits in its calculation capacity. That means that any p-value smaller than 1e-17 automatically rounds to 0.0. In this version, the function chi2.sf from scipy.stats.distributions is used to calculate very small p-values, but this function also has an upper limit (311 significant digits). For chi-square values greater than 1424 (df=1), the resulting p-value is rounded to 0.0. Significance levels then estimated at p-value < 1e-311 refer to values whose number of significant digits exceeds the calculation capacity of the most precise available functions. The primary objective of this test is to determine heterogeneity; the lack of precision at these low p-values is not problematic.

