## Supplemental Figures and Tables for "Genome-Wide Search for Candidate Drivers of Adaptation Reveals Genes Enriched for Shifts in Purifying Selection (SPurS)"

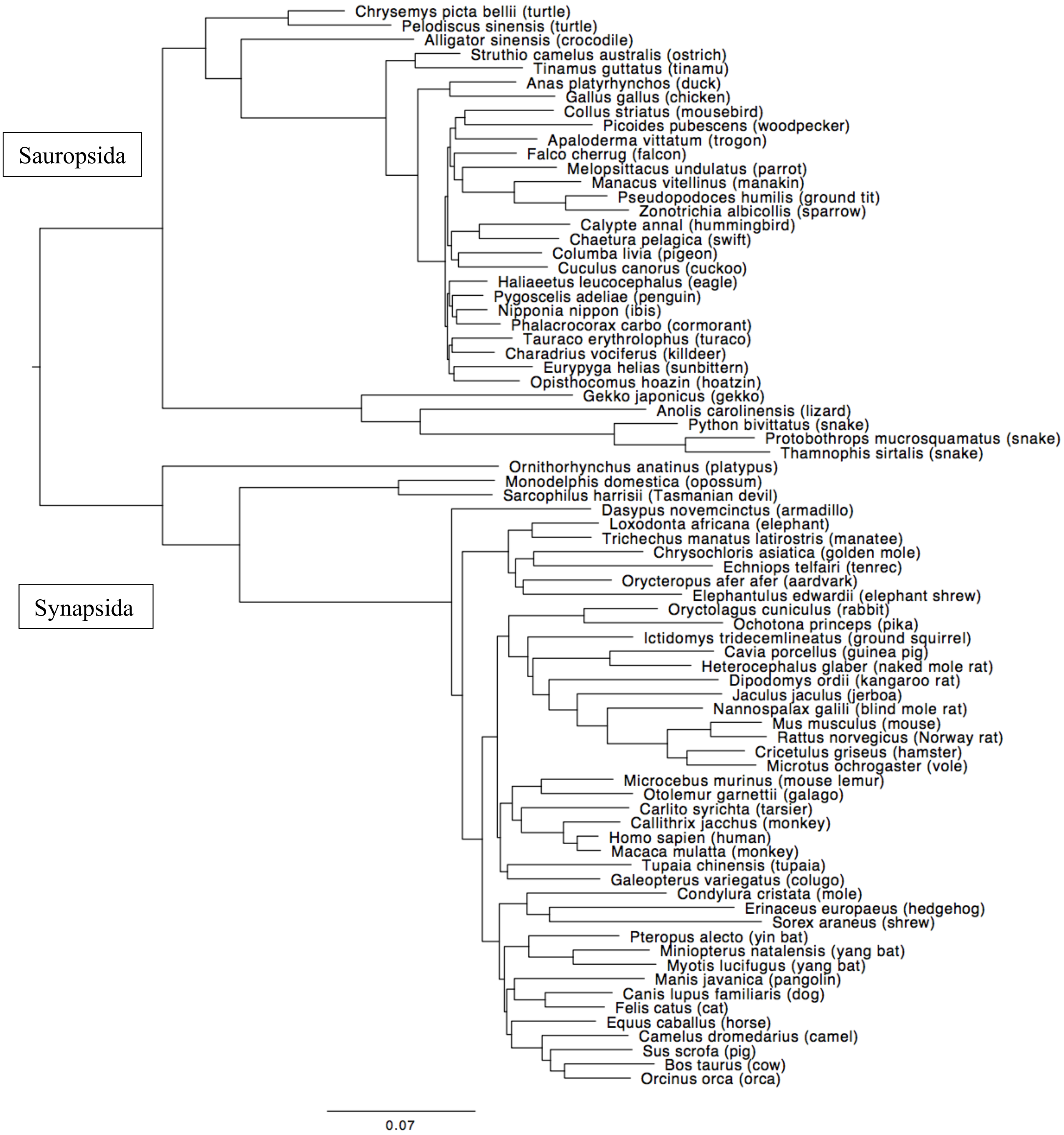


**Figure S-1**. Species phylogeny for all taxa included in SPurS analysis. Unrooted maximum likelihood protein tree representing the phylogeny of Sauropsids and Synapsids, in our analysis, constructed from concatenated single copy coding exons sampled across the genome. The tree was built using PhyML (LG model, 6 gamma-distributed rate classes, SPR search) from an alignment with an average of 90,457 residues per species (minimum 77,051). Except for the well accepted monophyly of Sauropsida and Synapsida, the precise topology of this tree is unimportant for this work and branch confidences were not computed. Armadillo is most likely misplaced on the tree; it should be sister to the elephant clade (doi: 10.1093/gbe/evv261) and the branch order near the base of Neoaves is very hard to resolve and remains controversial (doi: 10.1126/science.1253451 and doi: 10.1038/nature15697).

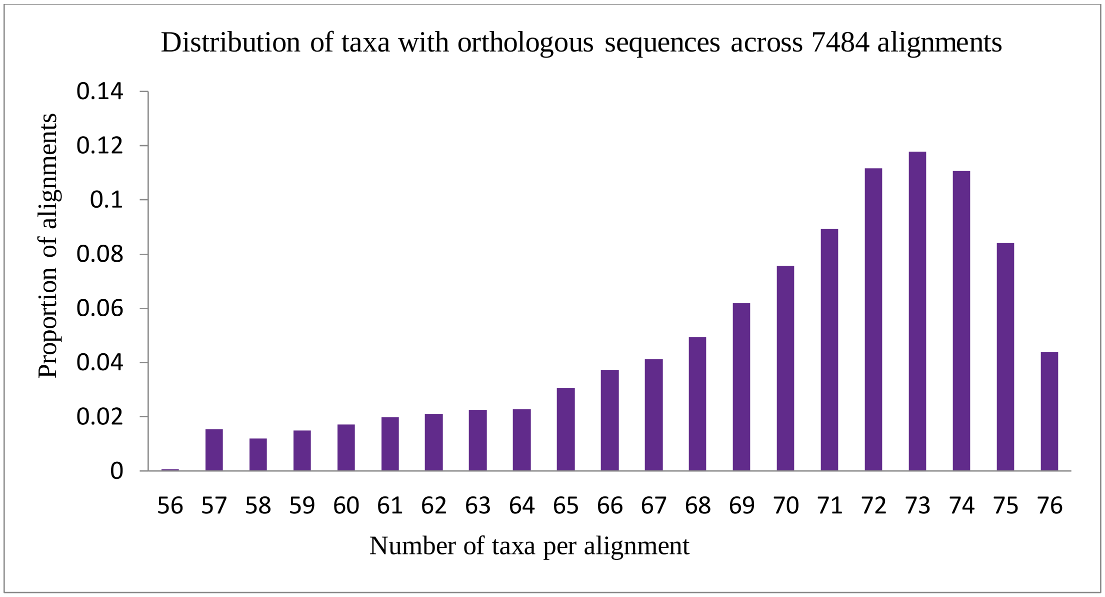


**Figure S-2.** Taxa with orthologous protein sequences, by percent of alignments. The number of species with orthologous sequences for each gene is represented on the x-axis, and the proportion of alignments with each number of species represented is on the y-axis. The absence of genes in certain species and lineages may be due to incomplete genome assembly, incorrect prediction, or true gene loss.

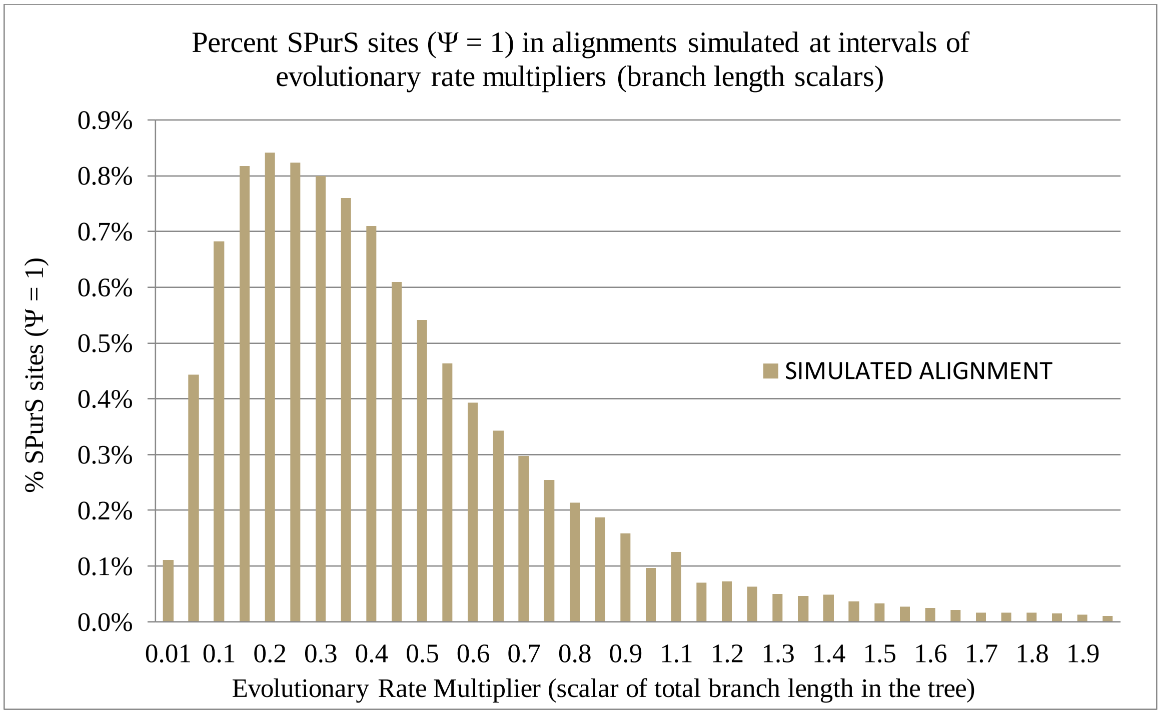


**Figure S-3.** Simulated alignments generated to determine the rate multiplier that maximizes observed complete SPurS sites. Multiple-species alignments were simulated in Seq-Gen, conditioned on the species phylogeny, and each one conditioned on varying rate multipliers.

**Table S-1.** Species Sampled for SPurS Analysis. Scientific and common names are provided, as well as taxonomic classifications. Species with the classifications Eutheria, Metatheria, and Monotremata were included in the "Synapsida" group; all others are included in the "Sauropsida" group. Species code names were used to name the sequences in each alignment file; scientific names are used for formal classification, and common names describe each species colloquially. Taxonomic classifications vary.

| Species Code Name | Scientific Name | Common Name | Superclass/ Class | Subclass/ Superorder/ Order/Suborder | Superfamily/ Family/Subfamily |
|---|---|---|---|---|---|
| Aplat | Anas platyrhynchos | duck | Galloanserae | Anseriformes | Anatidae |
| Ggal | Gallus gallus | chicken | Galloanserae | Galliformes | Phasianidae |
| Chavoc | Charadrius vociferus | killdeer | Neoaves | Charadriiformes | Charadriidae |
| Clivi | Columba livia | pigeon | Neoaves | Columbiformes | Columbidae |
| Cuccan | Cuculus canorus | cuckoo | Neoaves | Cuculiformes | Cuculidae |
| Tauery | Tauraco erythrolophus | turaco | Neoaves | Cuculiformes | Musophagidae |
| Eurhel | Eurypyga helias | sunbittern | Neoaves | Eurypygiformes | Eurypygidae |
| Fcher | Falco cherrug | falcon | Neoaves | Falconiformes | Falconidae |
| Opihoa | Opisthocomus hoazin | hoatzin | Neoaves | Opisthocomiformes | Opisthocomidae |
| Zalb | Zonotrichia albicollis | sparrow | Neoaves/A.1 | Passeriformes | Emberzidae |
| Phumi | Pseudopodoces humilis | ground tit | Neoaves/A.1 | Passeriformes | Paridae |
| Manvit | Manacus vitellinus | manakin | Neoaves/A.1 | Passeriformes | Pipridae |
| Mundu | Melopsittacus undulatus | parrot | Neoaves/A.1 | Psittaciformes | Psittaculidae |
| Halleu | Haliaeetus leucocephalus | eagle | Neoaves/A.2 | Accipitriformes | Accipitridae |
| Colstr | Colius striatus | mousebird | Neoaves/A.2 | Coliiformes | Coliidae |
| Picpub | Picoides pubescens | woodpecker | Neoaves/A.2 | Piciformes | Picidae |
| Apavit | Apaloderma vittatum | trogon | Neoaves/A.2 | Trogoniformes | Trogonidae |
| Nipnip | Nipponia nippon | ibis | Neoaves/B | Ciconiiformes | Threskiornithidae |
| Egrgar | Egretta garzetta | egret | Neoaves/B | Pelecaniformes | Ardeidae |
| Padel | Pygoscelis adeliae | penguin | Neoaves/B | Sphenisciformes | Speniscidae |
| Chapel | Chaetura pelagica | swift | Neoaves/C | Apodiformes | Apodidae |
| Calann | Calypte anna | hummingbird | Neoaves/C | Apodiformes | Trochilidae |
| Strcam | Struthio camelus australis | ostrich | Paleognathae | Struthioniformes | Struthionidae |
| Tingut | Tinamus guttatus | tinamu | Paleognathae | Tinamiformes | Tinamidae |
| Alsin | Alligator sinensis | crododilian | Sauropsida | Crocodilia | |
| Chasi | Chrysochloris asiatica | golden mole | Eutheria | Afrotheria | Chrysochloridae |
| Lafr | Loxodonta africana | elephant | Eutheria | Afrotheria | Elephantidae |
| Eedw | Elephantulus edwardii | elephant shrew | Eutheria | Afrotheria | Macroscelididae |
| Oafe | Orycteropus afer afer | aardvark | Eutheria | Afrotheria | Orycteropodidae |
| Etel | Echinops telfairi | tenrec | Eutheria | Afrotheria | Tenrecidae |
| Tman | Trichechus manatus latirostris | manatee | Eutheria | Afrotheria | Trichechidae |
| Cfam | Canis lupus familiaris | dog | Eutheria | Carnivora | Caniformia/Canidae |
| Fcat | Felis catus | cat | Eutheria | Carnivora | Feliformia/Felidae |
| Btau | Bos taurus | cow | Eutheria | Cetartiodactyla | Bovidae |
| Cbact | Camelus bactrianus | camel | Eutheria | Cetartiodactyla | Camelidae |
| Oorc | Orcinus orca | orca | Eutheria | Cetartiodactyla | Cetacea/Odontoceti |
| Sscr | Sus scrofa | pig | Eutheria | Cetartiodactyla | Suidae |
| Minnat | Miniopterus natalensis | yang bat | Eutheria | Chiroptera/ Yangochiroptera | Vespertilionidae |

| | | | | | |
|---|---|---|---|---|---|
| Mluc | Myotis lucifugus | yang bat | Eutheria | Chiroptera/ Yangochiroptera | Vespertilionidae |
| Palec | Pteropus alecto | yin bat | Eutheria | Chiroptera/ Yinpterochiroptera | Pteropodidae |
| Galvar | Galeopterus variegatus | colugo | Eutheria | Dermoptera | Cynocephalidae |
| Eeur | Erinaceus europaeus | hedgehog | Eutheria | Eulipotyphla | Erinaceidae |
| Sara | Sorex araneus | shrew | Eutheria | Eulipotyphla | Soricidae |
| Ccri | Condylura cristata | mole | Eutheria | Eulipotyphla | Talpidae |
| Ocun | Oryctolagus cuniculus | rabbit | Eutheria | Lagomorpha | Leporidae |
| Opri | Ochotona princeps | pika | Eutheria | Lagomorpha | Ochotonidae |
| Ecab | Equus caballus | horse | Eutheria | Perissodactyla | Equidae |
| Mjav | Manis javanica | pangolin | Eutheria | Pholidota | Manidae |
| Cjac | Callithrix jacchus | NW monkey | Eutheria | Primate/Haplorhini | Callitrichidae |
| Mmul | Macaca mulatta | OW monkey | Eutheria | Primate/Haplorhini | Cercopithecidae/ Cercopithecinae |
| Hsap | Homo sapiens | human | Eutheria | Primate/Haplorhini | Hominidae |
| Tsyr | Carlito syrichta | tarsier | Eutheria | Primate/Haplorhini | Tarsiidae |
| Mmur | Microcebus murinus | mouse lemur | Eutheria | Primate/Strepsirrhini | Cheirogaleidae |
| Ogar | Otolemur garnettii | galago | Eutheria | Primate/Strepsirrhini | Galagidae |
| Hgla | Heterocephalus glaber | naked mole rat | Eutheria | Rodentia | Bathyergidae |
| Cpor | Cavia porcellus | guinea pig | Eutheria | Rodentia | Caviomorpha/Caviidae |
| Jjac | Jaculus jaculus | jerboa | Eutheria | Rodentia | Dipodidae |
| Dord | Dipodomys ordii | kangaroo rat | Eutheria | Rodentia | Heteromyidae |
| Cgri | Cricetulus griseus | hamster | Eutheria | Rodentia | Muroidea/Cricetidae |
| Moch | Microtus ochrogaster | vole | Eutheria | Rodentia | Muroidea/Cricetidae |
| Mmus | Mus musculus | mouse | Eutheria | Rodentia | Muroidea/Muridae |
| Rnor | Rattus norvegicus | Norway rat | Eutheria | Rodentia | Muroidea/Muridae |
| Nangal | Nannospalax galili | blind mole rat | Eutheria | Rodentia | Muroidea/Spalacidae |
| Stri | Ictidomys tridecemlineatus | ground squirrel | Eutheria | Rodentia | Sciuridae |
| Tbelc | Tupaia chinensis | tupaia | Eutheria | Scandentia | Tupaiidae |
| Dnov | Dasypus novemcinctus | armadillo | Eutheria | Xenarthra | Dasypodidae |
| Shar | Sarcophilus harrisii | Tasmanian devil | Metatheria | Dasyuridae | Dasyuridae |
| Mdom | Monodelphis domestica | opossum | Metatheria | Didelphimorphia | Didelphidae |
| Oana | Ornithorhynchus anatinus | platypus | Monotremata | Monotremata | Ornithorhynchidae |
| Gekjap | Gekko japonicus | gekko | Sauropsida | Squamata/ Bifurcata/Gekkota | Gekkonidae |
| Acar | Anolis carolinensis | lizard | Sauropsida | Squamata/Iguania | Dactyloidae |
| Thasir | Thamnophis sirtalis | snake | Sauropsida | Squamata/ Serpentes | Colubridae |
| Pmol | Python bivittatus | snake | Sauropsida | Squamata/ Serpentes | Pythonidae |
| Promuc | Protobothrops mucrosquamatus | snake | Sauropsida | Squamata/ Serpentes | Viperidae |
| Cpic | Chrysemys picta bellii | turtle | Sauropsida | Testudines/ Cryptodira | Emydidae |
| Pesin | Pelodiscus sinensis | turtle | Sauropsida | Testudines/ Cryptodira | Trionychidae |
